# CalFluxTools: An R package for analysis of high-throughput calcium oscillation screening data

**DOI:** 10.64898/2026.09.15.751806

**Authors:** Andrew C. Patt, William F. Borschel, John Braisted, Danyal Raza, Chia-Kuei Wu, Atena Farkhondeh, Caroline Strong, Jiajing Zhang, Emily Lee, Bryan J. Traynor, Ewy A. Mathé

## Abstract

Whole plate detection of calcium flux signals using fluorescence is a powerful technology for monitoring oscillatory activity of excitable cells in a high throughput fashion that sees broad applications in neurotoxicity and cardiotoxicity research. However, analysis of calcium oscillation profiles can output more than 60 peak kinetic parameters, which makes manual data analysis challenging. We have created the CalFluxTools R package, an automated pipeline for parsing plate data for 384 well plates, generating quality control metrics and plots, and performing data analysis, including T-tests, Z-score analysis, and machine learning prediction of compound toxicity values. CalFluxTools requires a plate map and experimental manifest that includes parameter filtration specifications, as well as data in the “.statall” format. CalFluxTools runs in less than a minute on a Macbook Pro with a 2.6 GHz 6-Core Intel Core i7 processor.

## INTRODUCTION

As the cost and time of developing new therapeutic agents increases, monitoring of intracellular ion dynamics provides a crucial means for drug discovery and assessing experimental agents for toxic effects. High-throughput (HT), whole plate kinetic imaging readers that can simultaneously measure fluorescent emissions across an entire microplate (96-, 384-, and 1536-well plate formats) to measure real-time calcium oscillations have enabled applications in a diverse range of high throughput assay settings including drug repurposing and toxicity screening^1,2^. A typical assay may utilize cells preloaded with fluorescent dye and differences in fluorescence emission coinciding with compound treatment results as intracellular ion concentrations specific to the dye change, for which both the Molecular Devices Fluorometric Imaging Plate Reader (FLIPR) and the Hamamatsu Functional Drug Screening System (FDSS) platforms are amenable.

The flexibility of these platforms can accommodate numerous assay and experimental designs, cellular models, and reporter systems. Screening assays for measuring excitable cell activity are often designed to record calcium flux from non-excitable cells before and during or after compound treatment^3–6^. This assay design often limits the number of the endpoint measurements to the minimum or maximum response during dosing or the difference in steady-state intensity before and after treatment.

Excitable cells, including neurons and cardiomyocytes, have been valuable for identifying compound arrhythmia and seizure liability in vitro^7–10^. While using neurons and cardiomyocytes can provide a more physiological screening assay, intrinsic calcium oscillations typically seen in excitable cell types increases the calcium signal complexity and the number of measurable endpoint parameters^11,12^. As an example, PeakPro 2 analysis of raw calcium oscillations can return over 30 peak kinetic properties, outputting up to 63 different endpoint measurements, including 9 variations of peak width. This large number of measurements combined with tens of parameters describing redundant characteristics of calcium peak kinetics makes analysis of HT calcium oscillation data challenging as existing methods for analyzing this data rely on custom scripts and proprietary software.

In order to overcome these challenges, we have implemented the open-source CalFluxTools R package, which automates analysis of peak kinetics data. The current implementation described here was developed and validated using .statAll output generated by ScreenWorks Peak Pro 2 (Molecular Devices); compatibility with other kinetic plate-reader platforms has not been evaluated. CalFluxTools automatically parses these files and generates a series of visuals and analyses for quality control and toxicity prediction purposes. The standardized outputs are organized into output directories specified by the user and tailored to the experimental design. Outputs of the package allow users to inspect data for quality concerns on a per-well basis, compare results across plates, timepoints, and concentrations, and identify toxic exposures. Excel input sheets allow users to finely tune their analysis with custom well masking, set replacement values for missing peak parameter measurements, and meta-data entry that makes the package useable for programmers and non-programmers alike. In all, CalFluxTools greatly simplifies and streamlines the analysis of calcium oscillation peak kinetics data, improving efficiency and reproducibility.

## MATERIALS AND METHODS

### Expected Input Data

CalFluxTools is currently designed to accommodate high-throughput screening experiments in a 384-well plate format. The software can accommodate a variety of cellular models, compounds at various doses for dose-response analysis, and time points (**Fig.1**). The package divides experimental designs into two major categories, paired and unpaired (**Supplemental Fig. 1**). A paired analysis is a special experimental design wherein an experimental plate is “paired” with a time point 0 reference plate, which controls for inter-well variability. In a paired analysis, the experimental data for statistical testing and machine learning analysis are transformed relative to the reference plate (see below). In contrast, an unpaired design simply uses the experimental values post filtration and imputation as the values for analysis. Paired experimental design allows for the generation of several additional outputs, including an excel file of the transformed data, a heatmap of coverage relative to the reference plate, and T-testing for parameter shifts relative to the reference plate, highlighting parameters with the largest deviation from the reference condition.

**Figure 1:**
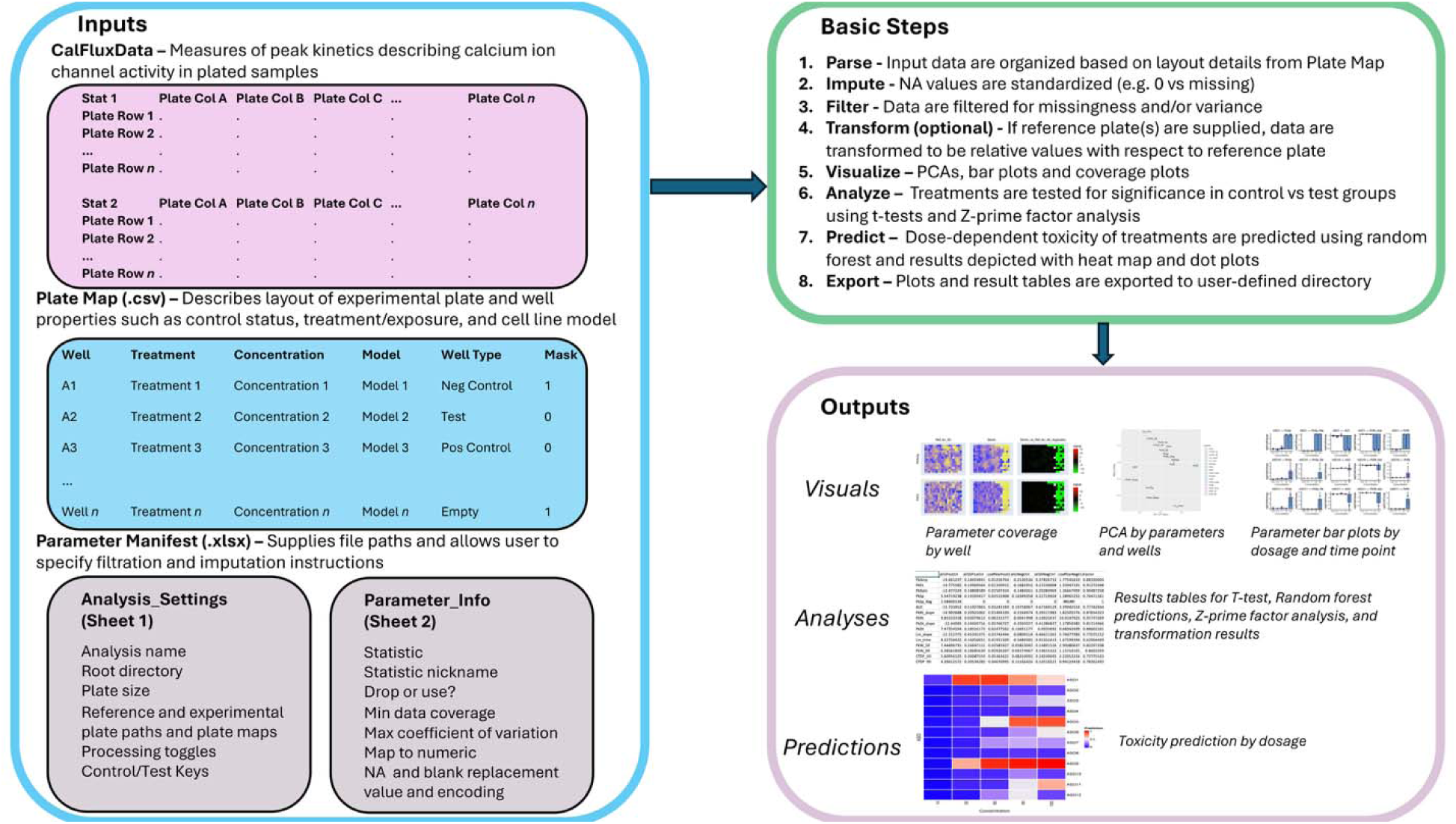
CalFluxTools Schematic. CalFluxTools is an R package for automated parsing and analysis of calcium flux data. The package accepts three inputs: calcium flux kinetics data in the .statALL format, as output by the PeakPro software, a plate layout manifest referred to as a plate map, and a parameter manifest. The .statALL data file is a format from the proprietary PeakPro software that organizes peak parameters into repeating dataframes that contain individual parameters in a layout mimicking the original plate layout. CalFluxTools parses this information and assembles it into the “CalFluxData” S4 class. The Plate Map is a csv table that lists the well numbers for the experimental 384-well plate, and includes metainformation such as treatment(e.g. compound/exposure)

### Package Description

CalFluxTools is run using a single wrapper function, *processCalFluxData*, which reads information from a user-supplied manifest to inform all downstream functions. StatAll files commonly contain 63 parameters that describe various aspects of calcium ion channel peak kinetics. A basic visual describing these parameters is provided in **Supplemental Fig. 2**. Because parameters are described with different units, with different considerations for missingness and zero values, a generic S4 class was constructed to store all data and meta-data associated with statAll data, called the *CalFluxData class. CalFluxData* objects typically contain objects from more specialized S4 classes, *plateSet* and *plate*. *plateSet* is a special class for experimental designs that incorporate a reference plate (described below), which *processCalFluxData* recognizes as a “paired” analysis. The *plate* class is the generic class for storing many facets of plate-related data, including readings by parameter, information about the plate layout, and information regarding plate data processing, including transformations and imputations.

The manifest used to direct *processCalFluxData* is a user-supplied file in the .xlsx format. The manifest serves multiple purposes. First, it contains the locations of the statAll data files as well as the plate maps corresponding to the data. The plate maps are another standardized format that must be filled out by the user. The plate map encodes metadata for each of the wells in the supplied experimental plate, including compound, concentration, model type, well type (e.g. Test, Positive Control, Negative Control, Empty) and mask (should the well be dropped from analysis). Second, the manifest includes specifications for how the analysis should be run, including plate pairings (if applicable), data transformation method, and export locations. Third, the manifest contains parameter-specific information designated by the user, including thresholds for filtration based on coverage or coefficient of variation, strategies to replace blank vs NA values, and manual use/drop designation. In this way CalFluxTools enables very granular control over parameter quality that are customizable based on the quality standards desired by the user.

Once inputs are assembled, CalFluxTools analysis proceeds as outlined in **Figure 1**. The package uses custom parsers to read in statAll files and assemble the data into corresponding slots in the *plate* class. Next, data are imputed based on the specifications from the parameter manifest, wherein instructions are encoded for handling missingness in a parameter-specific manner. This is because NA and blank values can have different meanings based on the parameter they correspond to and must be handled appropriately. Next data are filtered for missingness or for high variance (as determined by coefficient of variation). Again, appropriate thresholds are designated per parameter by the user in the manifest. Next, in the case of a paired analysis, a data transformation is optionally performed. The transformed values are computed as the log base-two fold-change between the experimental and the reference plate. The transformed data are exported to the export directory if the user wishes to analyze them in an alternate external program.

Following parsing and processing of the data, several visualizations are generated for quality control and analysis. All visuals (along with all other output files) are exported to the directory specified in the parameter manifest. First, CalFluxTools outputs Principal Component Analysis (PCA) plots, with respect to both parameters and wells. PCAs that display wells are labeled with control-level designations, and a special PCA plot filtered down to just control wells is also generated. A PCA of the parameters in only control wells is provided as well. Next, CalFluxTools outputs heatmaps that depict coverage by parameter in each well, alongside the relative shift from the refence plate, if applicable. These heatmaps can be used to identify spatial patterns in missingness when they occur. The package also generates bar plots of each parameter by compound dosage and time point, for easy synthesis and inspection of results.

Several output data files are also generated that include processed data or analytical results as well. First, an excel spreadsheet color-coded with Z scores by-well is output, which also aids in identifying spatial anomalies in experimental plates. Next, Z prime factors are computed for each parameter for each time point. Z prime factors are calculated as in **Equation 1**, where *σ_p_* and *σ_n_* are the standard deviation across all positive and negative control readings, respectively, and *µ_p_* and *µ_n_* are the mean across all positive and negative control readings, respectively.

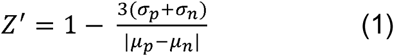

Z prime factors are used to assess separation from positive and negative control compounds, to ensure high quality results^13^. The software also exports the results of T-testing between the experimental and reference plates for each time point and dosage combination.

Lastly, CalFluxTools uses the RandomForest^14^ approach generate continuous toxicity predictions in the supplied exposures, trained on readings from known toxic (positive control) and nontoxic (negative control) exposures. The randomForest algorithm is run with an *mtry* equaling a third of the number of input variables, and an *ntree* of 1000, trained with 10-fold cross-validation. Controls are specified as toxic or nontoxic with a toxicity of value of 1 and 0, respectively, and continuous predictions are generated to produce a toxicity score between 0 and 1 for test wells. The results of the machine learning predictions are output in excel tables, dot plots, and heatmaps, that summarize the results across all dosages, time points and exposures. A benchmark dose is automatically calculated from the RandomForest toxicity predictions representing the lowest tested concentration at which a compound’s average predicted toxicity score is greater than 0.5. A variable importance plot is also output so that users can assess which parameters contributed most to the predictions made by the algorithm. Variable importance is assessed using the varImp function from the *caret* package^15^, and represents a measure of the increase of Mean Squared Error in predictions if a variable’s values are permuted.

Two example data sets along with their corresponding manifests are included in the *gh-pages* branch of the package github repo. To demonstrate use of the package, a vignette called *CalFluxTools_example_data.Rmd* provides an example analysis of the packaged data. This vignette shows the use of the high-level wrapper function, *processCalFluxData*, as well as several helper functions provided by the package. The example manifests are particularly helpful in providing a template for a properly formatted input that is readable by the package.

### Test-Case #1 –Simulated Data

ScreenWorks5.1 (Molecular Devices) was used to output a statAll file containing all 63 peak parameters with the PeakPro2 module using previously obtained FLIPR data in 384-well format. The file was edited in Excel (Microsoft Office) by manually deleting all corresponding parameter values from all wells and replacing each parameter with a single arbitrary value that varied between parameters. SD parameters were typically set to 10% of the corresponding mean parameter value and parameters relating to early afterdepolarization (EAD)-like events were set to either “0” or “N/A” and used as the “baseline” FLIPR response. A copy of the baseline statAll file was then used to generate post-dosing data by manually altering parameter values in Excel.

22 parameters were arbitrarily chosen to be altered to simulate dosing of vehicle (negative control), a toxic control (positive control), and a compound with high toxic potential (Toxic#1) and a compound with low toxic potential (Non-Toxic#1) across four concentrations. Parameter values remained unchanged for wells designated as negative controls (n=6), while values for positive control wells (n=6) were altered to represent a ∼50 –75% decrease in activity. While values of most parameters were lowered to simulate toxicity induced inhibition, peak spacing was increased to correspond to a smaller peak count. Changes in both rise and decay kinetics were set to reflect a decrease in time values, but an increase in slope values.

Parameter values of wells assigned to dosing of Toxic#1 were typically altered starting at the lowest concentration. Analysis of responses containing a single or zero peaks results in blank or “N/A” output values for most parameters and were used for wells dosed at the highest concentration to simulate complete inhibition. Changes in parameter values for Non-Toxic#1 dosed wells typically started at the third highest concentration and values at the highest dose represented ∼25 –50% decrease in activity. Finally, Gaussian noise was added to each simulated parameter using a standard deviation equal to 0.1 * the standard deviation of values for that parameter across wells, with a minimum standard deviation value of 0.2.

All parameters except those relating to EAD-like events (Mean and SD EAD-like peak Rate, Number of EAD-like peaks per well, Mean and SD Number of EAD-like peaks) were included for analysis. Blank replacement values were individually set based on the directional change of the positive control values for that specific parameter. Paired analysis between the baseline and post-dosed simulated data was performed and this data set was used to assist with the development and testing of this R package.

### Test-Case #2 –Unpaired Analysis

A subset of Alzheimer’s disease (AD) phenotypic screening data previously published (REF) was examined using the unpaired analysis design. Prefrontal cortex (PFC) like spheroids from human iPSC-derived neurons containing WT or AD linked apolipoprotein e4/4 (APOE4) allele GABAergic neurons were dosed with clinically approved compounds for AD treatment (Donepezil [1 and 10uM], Memantine [10uM], Rivastigmine [10uM]) and compounds currently in pre-clinical development for AD treatment (EUK-134 [1 and 10uM], Hu-210 [10uM]). Vehicle (0.01% DMSO) control was dosed on both WT (n=4) and APOE4 (n=6) PFC-like spheroids. FLIPR recordings taken 90-minutes post-dosing were processed using the PeakPro2 module in ScreenWorks5.1 using positive event polarity, dynamic threshold of 5% RFU above baseline cutoff applied to all wells, and automated noise rejection applied to all wells. Wells were individually inspected for correct peak detection before outputting all 63 peak parameter measurements into a single statAll file.

All parameters except those relating to EAD-like events were included for analysis. Max CV was set to “200” for each parameter. Replacement values were restricted to Mean Peak Rate (“0”) and Peak Spacing (NA: “100”; Blank: “250”) parameters. WT spheroids dosed with DMSO were designated as the “Negative Control” and APOE4 spheroids dosed with DMSO were designated as “Positive Control” within the plate map file. Wells dosed with compound were set to “Test” in the plate map file and used for the prediction modeling.

### Test-Case #3 –Paired Analysis

Commercially purchased human iPSC-derived cortical spheroids (3D microBrain platform, StemoniX, Inc.) were used to screen for neuronal inhibition related to oligonucleotide-dependent acute neuronal toxicity. 384-well spheroid plates were maintained following the manufacturer’s instructions and used for screening after 14 days in culture. On the day of the experiment, the spheroid plate was pre-treated with a calcium dye for 2-hrs (FLIPR Calcium 6 Assay Kit, Cat#R8194, Molecular Devices) before a 5-minute baseline FLIPR recording was taken. Oligonucleotides with low, medium, or high in vivo acute neurotoxic potential were prepared in DPBS (without Ca2+) and dosed at 3 (n=6), 30 (n=6), 60 (n=6), 120 (n=4) uM. Vehicle treated wells were dosed with DPBS (n=21) and an oligonucleotide previously reported to cause acute neurotoxicity in vivo^16^ was dosed at 60 uM (n=6) and used as a toxic control. Oligonucleotide-dependent changes in activity were measured 90-minutes following dosing.

Baseline and 90-minute post-dosing FLIPR recordings were processed using the PeakPro2 module in ScreenWorks5.1. Peak detection was performed using positive event polarity, auto-assigned length for search vector length, and automated noise rejection applied to all wells. Peak detection was visually inspected for each well before outputting all 63 peak parameter measurements into a single statAll file.

All parameters except those relating to EAD-like events were included for analysis. Max CV was set to “200” for each parameter. Blank and NA replacement values were individually set based on the directional change of the positive control values for that specific parameter. Spheroids dosed with DPBS were designated as the “Negative Control” and spheroids dosed with the toxic control were designated as “Positive Control” within the plate map file. All other oligonucleotide dosed wells across all concentrations were set to “Test” in the plate map file and used for the prediction modeling.

## RESULTS

We validated the outputs of CalFluxTools using three FLIPR data sets that had been generated using the ScreenWorks software with the PeakPro2 module. These data sets were selected to test a variety of the functions of the package, encompass both experimental designs supported by the package, and test different exposure/types/biospecimens, as well as simulated data. CalFluxTools was able to automatically identify changes in peak kinetics that were intentionally added to altered data, as well as recapitulate known biology from two studies testing toxic/ameliorative treatments in 3-dimensional neuronal spheroid models.

### CalFluxTools captures engineered changes in manipulated FLIPR data

As described in the **Methods** section (Test-Case #1 –Simulated Data), a synthetically-generated statAll dataset was created in order to establish previously known “true positive” signals in test data to validate the ability of CalFluxTools to correctly classify unknown samples using a set of intra-plate positive and negative controls. 22 arbitrarily selected parameters were manipulated for these results. The data set comprised 6 “positive control” samples mimicking exposure to a toxic compound and 6 “negative control” samples mimicking exposure to vehicle dosing. “Low toxicity” and “High toxicity” exposures were simulated across 4 different dosages with 6 replicates apiece.

CalFluxTools automatically outputs several visuals and tables that aid in quality control for multiple aspects of FLIPR assays. We first examined variance in the time zero “Reference Plate” using the “Reference_Plate_QC” table automatically output by the package (**Fig 2A**). Unique to the paired analysis pipeline, the purpose of this table is to provide a quick numeric and visual reference for identifying outlier wells in the reference plate, which can inform well masking in future computational runs and alert the user to potential data quality issues. The table provides a z-score breakdown aggregated across all wells, as well as individual parameter breakdowns for more careful inspection.

**Figure 2:**
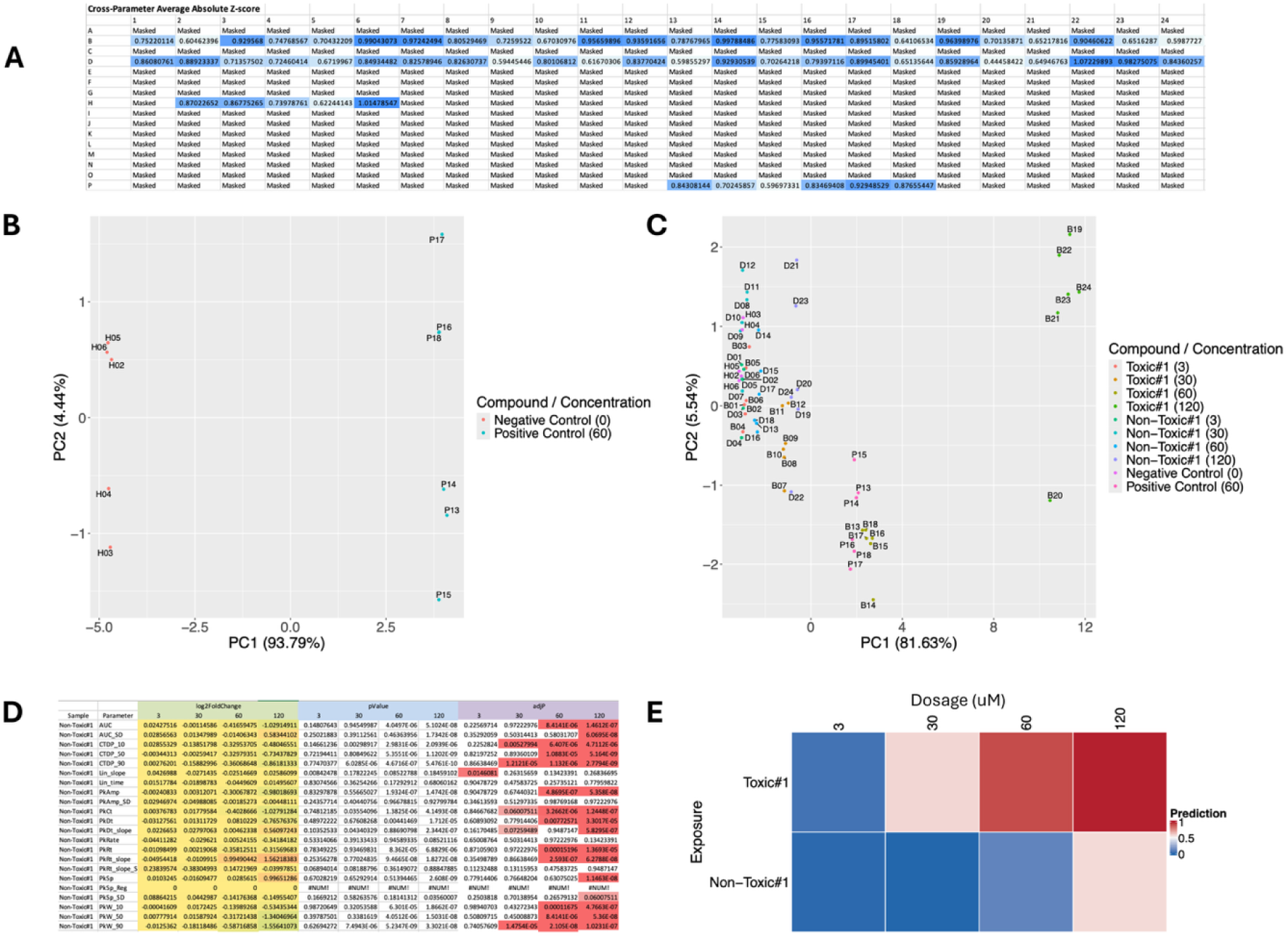
Example CalFluxTools outputs generated on simulated data capture expected trends using unsupervised clustering and supervised learning on FLIPR peak kinetic parameters. A) In “paired” analyses that include a timepoint zero reference plate, special outputs enable easy quality control of the reference plate, including this example table that highlights aggregate Z-score across parameters for each well for fast outlier identification B) PCA of control wells automatically output by the package shows clear separation of negative controls and positive controls in the test data. This feature allows for quick and easy assessment of control wells, making outlier identification easy and helping users ensure that negative and positive controls are sufficiently distinct. C) PCA of all wells demonstrates clear dose-dependent clustering between controls and different exposures. Higher doses of the simulated “Toxic” compound cause increasing shifts in the direction of the positive control cluster. D) To quickly identify exposures that induce significant change relative to the reference plate, and to identify the most impacted parameters, T-testing is automatically performed and color coded to highlight altered parameters post-exposure. E) The CalFluxTools machine learning module uses all FLIPR parameters to predict probabilities of class membership with respect to the positive and negative control wells. In this example visual, the simulated data has been correctly predicted to display a characteristic does-dependent toxicity response, with the highest dose of the “toxic” exposure displaying the closest relationship to the positive control wells.

Next, we examined the package’s PCA plot clustering samples in control wells using their values across all parameters (**Fig 2B**). As expected, the control well PCA displays good separation between the positive and negative control samples, indicating that controls are sufficiently distinct for downstream statistical testing and machine learning. Following this, we inspected the PCA plot encompassing all wells, including test wells (**Fig 2C**). Again, we found that the PCA plot captured anticipated patterns from the simulated data, with replicate wells assigned to the same treatment condition clustering closely together. Negative control wells remained clustered with non-toxic exposures and lower doses of the toxic exposure, whereas positive control wells grouped more closely with the maximum dosage of the toxic exposure. These findings show that CalFluxTools preserves the expected similarity structure of the data and can distinguish treatment classes based on multiparametric FLIPR phenotypes.

For statistical analysis/summarization of results, the CalFluxTools package provides several outputs. To get a granular breakdown of which exposures/dosages/parameters are significantly altered with respect to the reference plate, we examined the “T_test_results” table, which highlights parameters that are significantly different from their respective reference plate value following a paired T test and multiple test correction using the Benjamini-Hochberg method (**Fig 2D**). As expected we found both the higher doses of the toxic compound and the positive control compound significantly altered many parameters. Finally, CalFluxTools contains a machine learning module that uses the Random Forests approach to perform supervised learning and output a heatmap depicting probability of class membership with respect to the supplied control samples (**Fig 2E**). Our results demonstrated that CalFluxTools could stratify compound toxicity across dose levels. Samples corresponding to the toxic exposure were classified apart from vehicle-treated wells beginning at lower concentrations, consistent with the earlier and stronger parameter perturbations introduced into this group. In contrast, the low-toxicity condition remained closer to the negative-control profile at lower doses and separated primarily at the higher concentrations, matching the more modest simulated decrease in activity. Together, these results indicate that CalFluxTools not only groups biologically similar samples by unsupervised analysis, but also supports dose-responsive stratification of low- and high-toxicity phenotypes in downstream predictive modeling.

### CalFluxTools prioritizes compounds that restore wild-type function in mutant neuronal spheroid models

Next, to validate the predictions made by CalFluxTools, we reanalyzed data generated in a previous study by Strong et al.^17^ demonstrating that brain region-specific neural spheroids incorporating APOE4 GABAergic neurons exhibited peak kinetics characteristic of Alzheimer’s disease pathology (Test-Case #2 –Unpaired Analysis). The data further revealed that these changes could be partially rescued using clinically-approved therapies, resulting in kinetics that more closely resemble wild-type function, and that these changes could be detected using FLIPR data. The results of the previous study, which relied on a subset of manually selected FLIPR parameters, identified Donepezil and Memantine as leading compounds capable of restoring wild-type calcium activity patterns. We applied CalFluxTools to the same dataset to evaluate whether an automated, comprehensive analytical approach using all output FLIPR parameters could recapitulate and potentially enhance these findings without requiring prior parameter selection. All results for this analysis were generated in unpaired mode, as no reference plate was used for this data set.

We first examined the well-level PCA of the complete multiparametric dataset (**Fig 3A**). In general, we observed strong clustering based on exposure across all compounds and controls. Notably, wells treated with higher dosages of Donepezil and Memantine clustered preferentially with wild-type controls, while lower dosages and other treatments clustered more closely with the APOE4 disease model, mostly mirroring the phenotypic rescue identified through the targeted analysis approach of Strong et al. The consistency between our unbiased analytical approach and the original targeted analysis demonstrates that CalFluxTools maintains biological interpretability while reducing potential bias introduced by selective parameter analysis.

**Figure 3:**
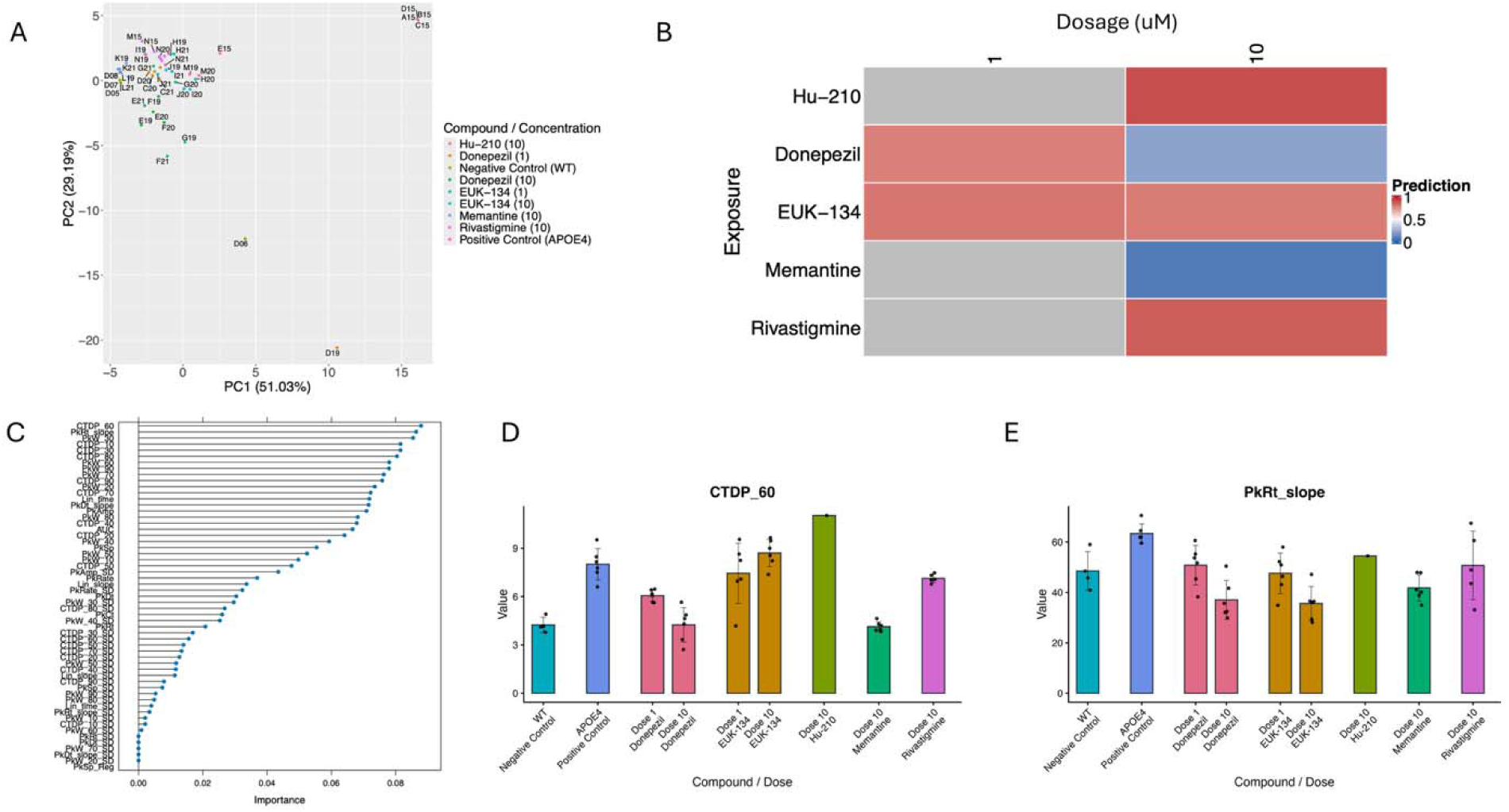
CalFluxTools prioritizes compounds restoring healthy function in mutant APOE4 neuronal spheroid Alzheimer’s model. A) Well–level PCA shows clinically approved treatments at higher doses cluster more readily with wild-type wells, while, other treatments cluster more closely with the mutant disease model. B) CalFluxTools machine learning module prioritizes Donepezil and Memantine as most restoring wild-type phenotype in mutant disease models, while the low dose of Donepezil and both doses of EUK-134 produced an intermediate phenotype. Bluer cells indicate a higher probability of wild type class membership, while redder cells indicate a higher probability of mutant class membership. C) Variable importance for machine learning model. CalFluxTools is able to incorporate all output variables into its predictions. D) Shifts observed in “calcium transient duration from peak to 60% height” (CTDP_60) parameter following compound exposure. CTDP_60 was the top predictive variable output by FLIPR. Values reflect transformation with regards to reference plate. E) Shifts observed in Decay Slope parameter following compound exposure. Values reflect transformation with regards to reference plate.

We next examined the outputs of the machine learning module of CalFluxTools. Consistent with the results of the original study, we found that wells treated with Memantine and Donepezil (both 10 μM) were predicted to be more likely to be belonging to the wild-type class (prediction value closer to 0), the lower dose of Donepezil and both doses of the experimental compound EUK-134 produced an intermediate classification, and all others aligned more closely with the mutant disease model class (prediction value closer to 1, **Fig 3B**). This finding further validated our results with respect to the original published data, as the lower Donepezil dose and both doses of EUK-134 were also identified as ameliorating the mutant phenotype when peak count and peak spacing were compared to control values. Next, examination of feature importance rankings from the random forest model revealed the parameters most discriminative for wild-type versus APOE4 classification (**Fig 3C**). Although in the original study, peak count and peak spacing were manually selected to illustrate the parameter shifts induced by the various tested compounds, our model prioritized calcium transient duration from peak to 60% height (CTDP_60) and rise slope (PkRt_slope) as the most discriminative parameters for predicting disease status (**Figs 3D** and **3E**). This finding demonstrates that CalFluxTools not only validates manually curated analytical approaches but may also reveal additional dimensions of drug-induced phenotypic modulation that could inform mechanism of action studies and improve prediction performance. Together, these results establish CalFluxTools as a robust platform for unbiased, comprehensive analysis of neurotoxicity responses in disease modeling and therapeutic screening applications.

### CalFluxTools accurately predicts neurotoxic exposures in neuronal spheroid models treated with oligonucleotides of previously characterized toxicity

Finally, we evaluated CalFluxTools using a paired-analysis dataset generated from human iPSC-derived cortical spheroids treated with oligonucleotides (ONTs) spanning previously characterized low, medium, and high acute neurotoxic potential (Test-Case #3 –Paired Analysis). Past studies using FLIPR to screen against ONT-dependent acute neurotoxicity have relied on datapoint counting with amplitude thresholds as the only end-point measurement and have not examined changes in peak kinetics. In this experiment, a 5-minute baseline FLIPR recording was collected before dosing and compared with a 90-minute post-treatment recording to serve as a reference plate, allowing the analysis to be performed in paired mode. Visual inspection of representative calcium traces showed that vehicle-treated spheroids retained spontaneous oscillatory activity, whereas ONT-treated wells displayed progressive disruption of calcium dynamics that matched known toxicity ranking of the compounds (**Fig. 4A**). In general, the low-toxicity and medium-toxicity ONTs produced relatively modest changes in activity, while the high-toxicity ONT induced pronounced declines in peak frequency, confirming our prior knowledge on the relative toxicity of the tested ONTs.

**Figure 4:**
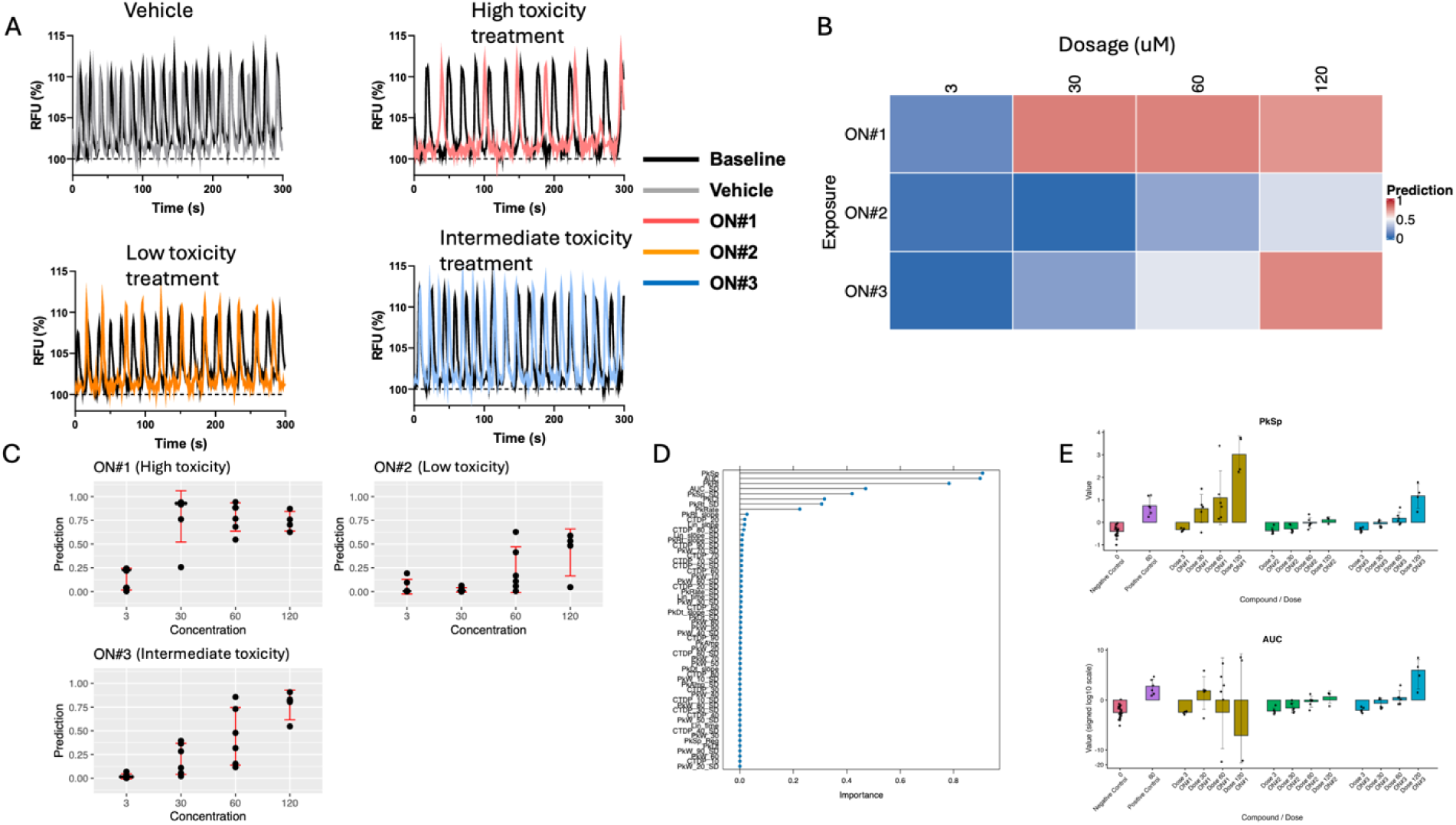
CalFluxTools correctly stratifies ONTs of previously known activities. A) Representative traces of calcium channel activity in treated spheroids show reduced activity in high toxicity ON#1, little activity reduction in the low toxicity ON#2, and moderate activity reduction in the intermediate toxicity ON#3. B) The CalFluxTools machine learning module successfully captured these trends in a dose-dependent fashion using output FLIPR parameters. C) Dot plots of CalFluxTools machine learning predictions resemble dose-response curves, and showcase the higher slope and peak of the highly toxic exposures compared to the intermediate and low toxicity exposures. D) Machine learning variable importance of the measured FLIPR parameters, which were highly distinct from the variables selected in the Alzheimer’s study. E) Barplots of most important parameters in machine learning model, Peak spacing and Area Under the Curve, post-transformation with regards to reference plate.

We next applied CalFluxTools to translate these observed peak kinetic shifts into quantitative toxicity predictions. Using vehicle-treated wells as negative controls and a previously established neurotoxic ONT as the positive control, the random forest model successfully stratified test ONTs in a dose-dependent manner (**Fig. 4B**). Heatmap outputs showed clear separation of exposures according to their expected toxicity, with prediction scores increasing from low- to medium- to high-toxicity ONTs as concentration increased. This pattern was recapitulated in the well-level dot plots, which resembled a dose-response relationship and demonstrated increasing predicted toxicity with increasing ONT dose (**Fig. 4C**). To better understand which peak-kinetic features drove this classification, we examined the highly prioritized variables from the machine learning model (**Fig. 4D**). Interestingly, the most important parameters were highly distinct from those obtained from the APOE4 AD data, underscoring the importance of unbiased and data-specific parameters for classifying samples (**Fig. 4E**). Together, these results show that CalFluxTools can accurately prioritize neurotoxic oligonucleotide exposures, recover expected dose-response relationships, and highlight the specific calcium-signaling features most associated with acute neuronal toxicity.

## DISCUSSION

To the best of our knowledge, CalFluxTools is the first open-source software tool specifically designed for analysis of 384-well high-throughput data generated on a high-thrpoughput, whole-plate reader kinetic imager platform. The package greatly simplifies a wide variety of tasks associated with data analysis, including missingness filtration, imputation, variable selection, data visualization and quality control. Because findings generated by the package are fast and reproducible, the software is useful for providing quality control feedback for intermediate experimental results, guiding further refinements and improvements to future iterations of experiments. The diverse array of QC plots and analytical results support a variety of different experimental designs, including multiple dosages, multiple timepoints, multiple exposures, and the presence and absence of a reference plate.

One of the most noteworthy features of CalFluxTools analysis is that it can leverage all parameters output by the FLIPR platform. Due to the high number of redundant, colinear parameters describing peak kinetics that are output by the platform, many analysts resort to manual selection of parameters to calculate statistics on, which introduces bias and potentially ignores the most informative parameters for a particular experiment. Alternatively, the RandomForest approach employed by the package considers each parameter when making predictions and builds predictors solely on the criteria of impact on model performance. This can improve the sensitivity of analyses over more targeted approaches and improve the ability of researchers to identify toxic/ameliorative compounds from FLIPR screening data.

Another major benefit of the CalFluxTools package is that it is highly flexible and scalable, being able to support measurements generated using a large number of exposures at different dosages and timepoints with several replicates. We anticipate that supporting the analysis of larger-scale assays will aid in unlocking the high-throughput potential of the FLIPR platform, which is more typically used for smaller-scale, targeted assays. Additionally, by providing the user the ability to manually filter and manipulate imputation guidelines for the package, we provide researchers with the ability to apply domain and study-specific knowledge to aid in neuro- and cardiotoxicity research and ensure that results are transparently generated and reproducible. CalFluxTools also produces useful intermediary files with post-transformation data in a convenient format that allows researchers to perform additional analyses in external programs that are not supported by the package currently.

Being a continuously upkept and actively developed software package, CalFluxTools holds promise for future expansions as well. We anticipate expanding the machine learning options available to users beyond RandomForests, in order to provide multiple options for optimizing prediction accuracy. We also plan to expand our options for supported experimental designs. First, we plan to add support for experimental plates with dimensions other than 16 rows x 24 columns. Second, we would like to improve support for comparing experimental results across plates. While currently separate plates are assumed to correspond with different timepoints, we recognize the utility of more flexible options with distributing samples across multiple plates. The use of a paired experimental setup is highly useful in this regard, as it can account for batch-effect concerns that may limit the use of multi-plate experimental designs.

## CONCLUSIONS

CalFluxTools provides a standardized, open-source framework for the analysis of high-throughput calcium oscillation-based peak-kinetics data that reduces the need for ad hoc scripting and manual parameter selection when performing statistical analysis and visualization. Across three distinct test cases, the package correctly identified engineered signal changes in simulated data, recapitulated known biology in a neuronal spheroid disease-model dataset, and accurately stratified oligonucleotides such as antisense oligonucleotides (ASOs) with previously characterized neurotoxic potential in a paired-analysis workflow. By integrating preprocessing, parameter-specific filtering and imputation, quality-control visualization, statistical testing, and machine learning-based toxicity prediction into a single pipeline, CalFluxTools enables rapid, reproducible, and interpretable analysis of complex calcium signaling phenotypes. We anticipate that this package will broaden access to comprehensive, high-throughput FLIPR data analysis and support more efficient neurotoxicity, cardiotoxicity, and phenotypic screening studies.

## Acknowledgements

This work was supported in part by the intramural program “Informatics Research” (ZIC TR000410) at the National Center for Advancing Translational Sciences, part of the National Institutes of Health, and National Institute on Aging (1ZIAAG000933). This research was supported in part by the Intramural Research Program of the National Institutes of Health (NIH). We would like to thank Dr. Donald Lo for enlightening discussions on FLIPR method development.

## Disclaimers

The contributions of the NIH author(s) were made as part of their official duties as NIH federal employees, are in compliance with agency policy requirements, and are considered Works of the United States Government. However, the findings and conclusions presented in this paper are those of the author(s) and do not necessarily reflect the views of the NIH or the U.S. Department of Health and Human Services.

