## Supplement for "CalFluxTools: An R package for analysis of high-throughput calcium oscillation screening data"

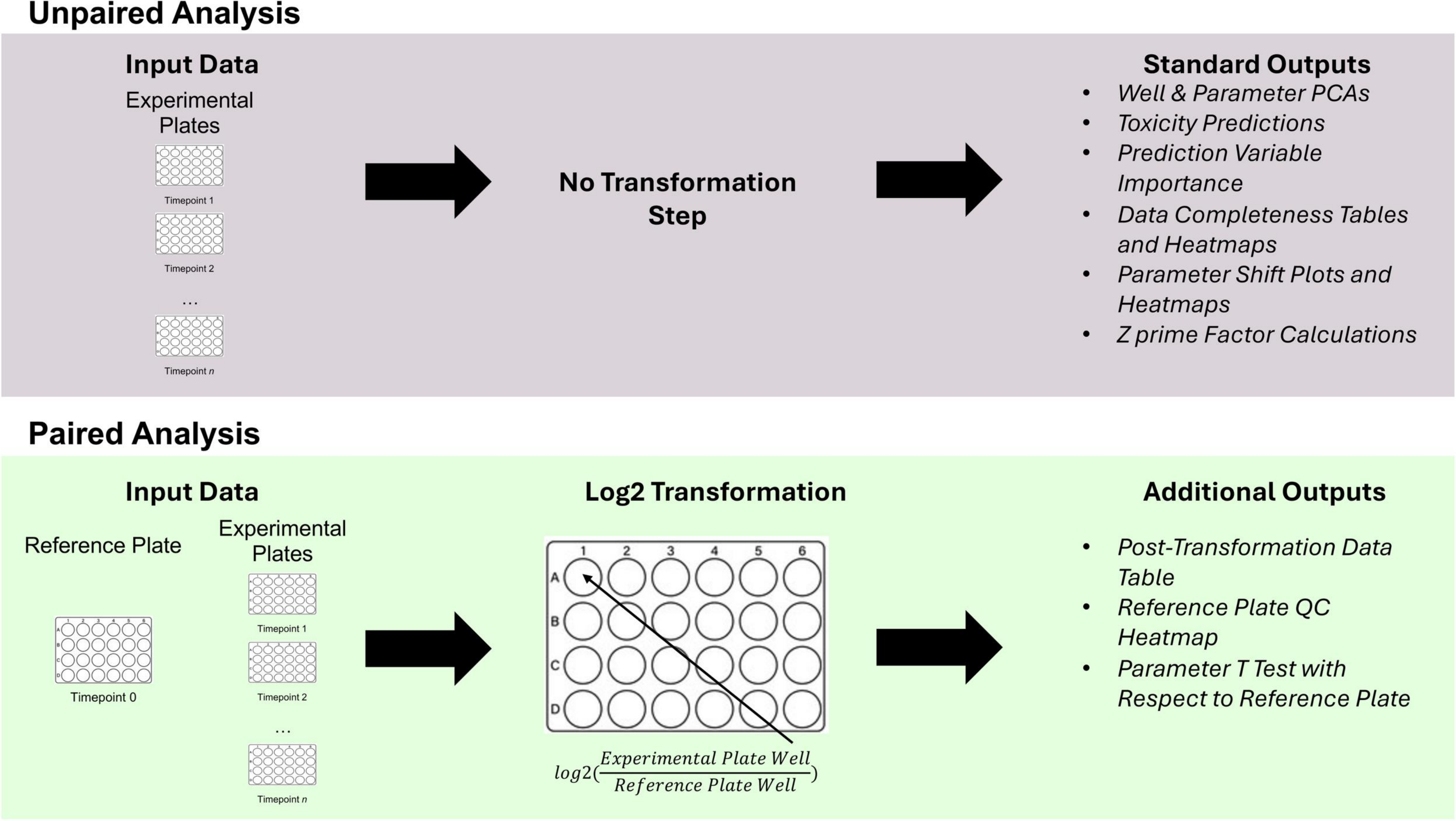


**Supplemental Figure 1**: Paired vs Unpaired analysis workflows. CalFluxTools accommodates two high-level experimental designs, depending on if a timepoint 0 reference plate was generated. In the case of an unpaired analysis, the plate values themselves are used for statistical testing and machine learning predictions. In contrast, a paired analysis

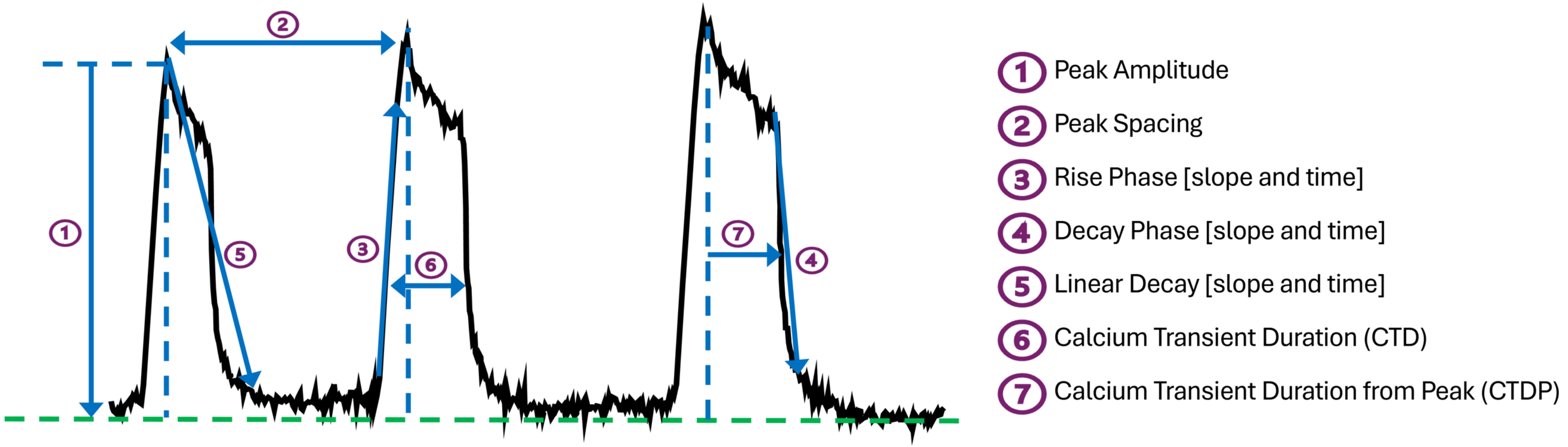


**Supplemental Figure 2**: Visual representation of peak kinetic parameters measurable with FLIPR. Calcium transient duration (CTD) parameters measure the peak width between 10 – 90% height from the peak while the calcium transient duration from peak (CTDP) parameters measure the width from the peak to 10 – 90% of the distance from the peak. Parameters relating to early afterdepolarization (EAD)-like events are not depicted.
